# At each site its diversity: DNA barcoding reveals remarkable earthworm diversity in neotropical rainforests of French Guiana

**DOI:** 10.1101/2020.09.21.307348

**Authors:** Marie-Eugénie Maggia, Thibaud Decaëns, Emmanuel Lapied, Lise Dupont, Virginie Roy, Heidy Schimann, Jérôme Orivel, Jérôme Murienne, Christopher Baraloto, Karl Cottenie, Dirk Steinke

## Abstract

Despite their recognized essential role in soil, earthworms in tropical environments are still understudied. The aim of this study was to re-evaluate the diversity at the regional scale, as well as to investigate the environmental and spatial drivers of earthworm communities. We sampled earthworm communities across a range of habitats at six locations in French Guiana using three different sampling methods. We generated 1675 DNA barcodes and combined them with data from a previous study. Together, all sequences clustered into 119 MOTUs which were used as proxy to assess species richness. Only two MOTUs were common between the six locations and 20.2 % were singletons, showing very high regional species richness and a high number of rare species. A canonical redundancy analysis was used to identify key drivers of the earthworm community composition. The RDA results and beta-diversity calculations both show strong species turnover and a strong spatial effect, resulting from dispersal limitations that are responsible for the current community composition. Sampling in different microhabitats allowed the discovery of 23 MOTUs that are exclusively found in decaying trunks and epiphytes, highlighting hidden diversity of earthworms outside of soil.

## 1. Introduction

Despite the fact that they host a large and complex array of species, soils still remain the most understudied habitat of terrestrial ecosystems (Bardgett and Putten, 2014; Decaëns, 2010; Wolters, 2001). Based on recent species richness estimates and because of a huge taxonomic deficit, soils are considered the third biotic frontier after tropical forest canopies and oceanic abysses (André et al., 1994; Giller et al., 1997; Wolters, 2001). Soil invertebrates in particular are key actors in most terrestrial ecosystems, including agroecosystems (Decaëns, 2010), as their activities are essential in sustaining key ecological processes (Lavelle et al., 2006). However, they remain insufficiently studied in comparison with other terrestrial aboveground organisms (Wolters, 2001). As a result, soil biodiversity patterns and their drivers remain largely unknown, especially at the global scale (Decaëns, 2010; Phillips et al., 2019). Several studies have already pointed out that soil organisms show different ecological patterns than those observed through the study of aboveground organisms (Cameron et al., 2019; Fierer et al., 2009; Phillips et al., 2019).

Earthworms are considered major faunal actors because of their importance in the maintenance of soil functions and the provisioning of soil ecosystem services (Lavelle et al., 2016). They have been characterized as soil engineers, due to their capacity at altering the soil structure, with important effects on its physical, chemical, and biological functioning (Jones et al., 1994; Lavelle et al., 2016, 2006). They are globally distributed, present in both temperate and tropical soils, with the exception of the driest and coldest regions of the planet, and can make up 60 % - 80 % of overall soil biomass (Amat et al., 2008). However, there are only a few studies so far focussing on earthworm community structure at a regional scale, and there is still a considerable lack of knowledge on their ecology and distribution particularly for tropical ecosystems (Feijoo, 2001; Fragoso, 1985; Jiménez, 1999; Lavelle, 1978).

One considerable roadblock to a better understanding of community ecology is the existence of a taxonomic impediment on soil biodiversity in general and earthworms in particular (André et al., 2001; Decaëns, 2010). Recent developments of molecular tools such as DNA barcoding have the potential to overcome the barriers of traditional taxonomy and thus facilitate the acquisition of new data that in turn can be used to describe the spatial distribution of species and communities in a rapid and comprehensive fashion. DNA barcoding uses the mitochondrial gene cytochrome c oxidase I (COI) as standard genetic marker for identification of animal species (Hebert et al., 2003). It can also be used as a mean to delineate Molecular Operational Taxonomic Units (MOTUs) in the absence of prior morphological identification. MOTUs are increasingly used to estimate taxonomic richness and to describe the spatial distribution of communities (Blaxter et al., 2005; Porco et al., 2013; Smith et al., 2005; Young et al., 2012). For instance, they were used to study diversity patterns of earthworm communities in the tropical rainforests of French Guiana (Decaëns et al., 2016). Authors were able to delimit 48 MOTUs that almost perfectly match with adult morphology, suggesting that MOTUs based on COI barcodes could in fact represent true biological species. The use of barcoding allows to take into account morphologically unidentifiable specimens such as juvenile earthworms or cocoons, as well as cryptic species, unlike the traditional taxonomy identification method. Unfortunately, the use of barcoding is still limited in the study of tropical earthworm communities, biasing the current datasets in tropical regions. With that, the classic Tropical Soil Biology and Fertility (TSBF) quantitative sampling method (Anderson and Ingram, 1989) that has been widely used in tropical studies to characterize earthworm biodiversity, but also other key members of soil biota, does not seem adapted to the context of tropical rainforests. For instance, Bartz et al. (2014) showed that more species were collected using the qualitative method and Decaëns et al. (2016) demonstrated that many species can be found in microhabitats others than the soil *sensu stricto*. Inadequate sampling methods may therefore generate a strong undersampling of earthworm species diversity in tropical ecosystems, and represent major barriers in the study and understanding of tropical earthworm communities in tropical regions, generating an underestimation of species diversity in the tropics.

For this study we generated a comprehensive data set at different spatial scales using samples collected during several expeditions in Amazonian tropical rainforests over the past few years. The sampling protocol coupled three methods comprising the traditional TSBF method associated with qualitative sampling in the soil and in other microhabitats. We wanted to analyse earthworm community patterns at regional, local, and habitat scale using newly generated DNA barcodes.

With the addition of new data for this region we expected an increase in the number of MOTUs at the regional scale (γ-diversity) and a low level of shared diversity between locations explained by a strong turnover (β-diversity); as it has been suggested that the high regional diversity of earthworms in the tropics could be the result of a high level of endemism and/or an higher beta-diversity towards the equator (Decaëns, 2010; Lavelle and Lapied, 2003). However, this pattern might not be observed at local or smaller scales (α-diversity). Indeed, some studies suggested a lack of local diversity peak in the tropics due to interspecific competition (Decaëns, 2010; Phillips et al., 2019), while others argued that a deficit of sampling, coupled with high levels of geographical turnover in community composition, could hide the existence of the latitudinal gradient (Lavelle and Lapied, 2003) or that there is no difference between tropical and temperate regions (Lavelle, 1983). In addition, we also expected that key environmental drivers such as soil properties (pH, organic carbon, soil texture etc.) or climate and habitat type (i.e. forest type) will influence earthworm diversity by shaping their community structure as it has been shown in previous studies (Mathieu and Davies, 2014; Phillips et al., 2019; Rutgers et al., 2016; Spurgeon et al., 2013).

## 2. Material and methods

### 2.1. Study sites

The sampling was performed in the Amazonian rainforests of French Guiana. This French overseas territory on the northern Atlantic coast of South America is 83,846 km^2^ which more than 95 % is covered by primary rainforest. It is characterized by a relatively uniform tropical humid climate, with only two seasons: a wet season between December and June, usually interrupted in February or March by a short drier period, and a dry season between July and November.

Four different locations, Galbao, Itoupé, Mitaraka and Trinité were sampled during rainy seasons between 2015 and 2019 (Figure 1) as part of the DIADEMA (DIssecting Amazonian Diversity by Enhancing a Multiple taxonomic-groups Approach) and DIAMOND (DIssecting And MONitoring amazonian Diversity) projects of the Labex CEBA (Center for the study of Biodiversity in Amazonia) and the expedition “Our Planet Reviewed” (Touroult et al., 2018). At each location, sampling was carried out in 11 to 19 1 ha plots, spaced by at least 500 m from each other, and representing contrasting local habitat types: hilltop (low canopy tropical rainforest located on somital position on shallow soils that dry quickly), plateau (high tropical rainforest on deep and well-drained soil), slope (tropical rainforest on slope and deep soils), swamp forests (tropical rainforest on hydromorphic soils) and specific vegetation formations of the inselbergs such as transition forests and rocky savannah when present (Supplementary Table 1).

- The Mont Galbao (lon / lat: −53.2830 / 3.5886) is a mountain range spreading over 6 km inside the National Amazonian Park of French Guiana (PAG) with peaks reaching 650 to 730 m above sea level (Figure 1). A total of 11 plots were sampled in January 2019, representing hilltop, slope, plateau and swamp forests at elevations ranging from 466 to 723 m.
- The Mont Itoupé (lon / lat: −53.0834 / 3.0222) is a mountain belonging to the Tabular Mountains chain located in the centre of the PAG (Figure 1). It is the second highest peak in the territory, with an altitude of 830 m above sea level, and it is composed of a large plateau covered by cloud forest. Sampling was conducted in January 2016, in a total of 14 plots including cloud forests above 800 m, and slope and plateau forests at lower elevation ranges (i.e. 440 to 635 m).
- The Mitaraka range was sampled in 2015 around a camp (lon / lat: −54.4503 / 2.2340) set up temporarily for this occasion (Figure 1). The massif is part of the Tumuc-Humac mountain chain located at the extreme South-West of French Guiana at the border to Brazil and Surinam. The landscape is classified as “high hills and mountains” (Guitet et al., 2013) and is characterized by the presence of massive inselbergs (isolated granite rock blocks that range several hundred meters above the lowland areas) and lowland forest. Sampling was conducted in 19 plots representing the complete range of habitats targeted for this study.
- The Trinité Natural Reserve was sampled around the Aya Camp (lon / lat: −53.4132 / 4.6024), which is located in the vicinity of a large isolated inselberg with an altitude of 501 m (Figure 1). Sampling took place in 2016 in 11 plots comprising swamp, slope and plateau forests, as well as hilltop and transition forests of the inselberg.
- Two locations were sampled in the Nouragues Natural Reserve in January and June 2011 around the two permanent research stations present in the area: Pararé station (lon / lat: −52.6730 / 4.0381) and Inselberg station (lon / lat: −52.6800 / 4.0883), both 6.4 km away from each other (Decaëns et al., 2016) (Figure 1). The former is located along the Arataye River, with vegetation dominated by lowland swampy forest, while the latter is situated at the foot of an isolated inselberg culminating at 411 m. A total of 41 plots were sampled, including inselberg habitats, plateau, slope and swamp forests (see Decaëns et al., 2016 for details).

**Figure 1:**
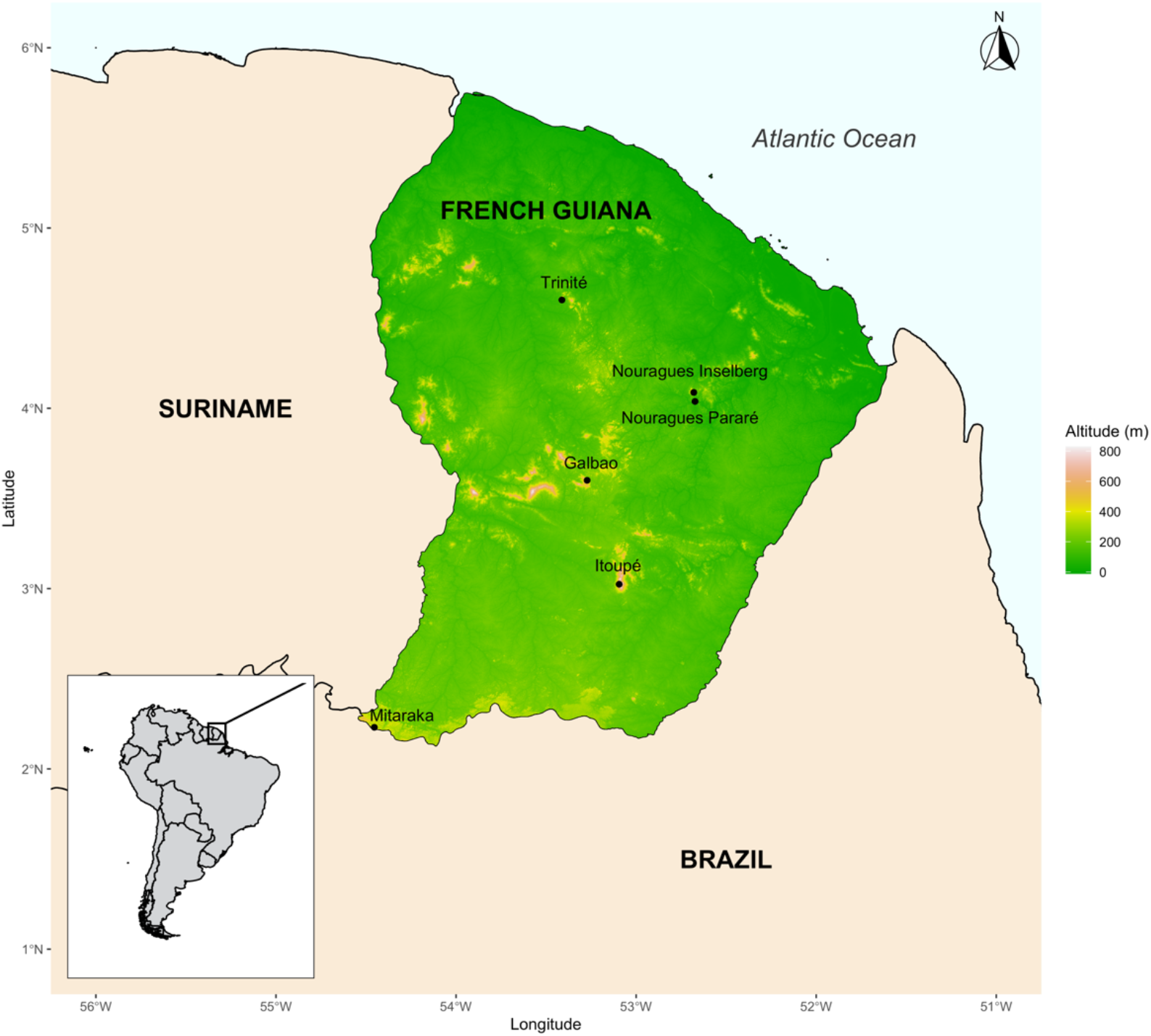
Map showing the six sampling locations in French Guiana. Data from Galbao, Itoupé, Mitaraka and Trinité were generated as part of this study, data from the Nouragues were taken from an earlier study (Decaëns et al., 2016).

### 2.2. Sampling (earthworms and soil)

In each 1 ha plot, earthworms were collected by combining three different approaches (protocol adapted from Decaëns et al., 2016): (1) Quantitative sampling (TSBF) by digging and hand-sorting three blocks of soil, each 25 x 25 x 20 cm (length x width x depth), located at the interior angles of a 20 m equilateral triangle in the center of the sampling plot; (2) Qualitative sampling by digging and hand-sorting an area of 1 m^2^ with a minimum depth of 20 cm, selecting an area with large earthworm casts (when available) within the sampling plot; (3) Micro-habitat sampling by visually inspecting all available micro-habitats (such as sandy to muddy sediments of stream banks, leaf litter accumulations, decaying trunks and epiphytic soils) for three researcher-hours (e.g. one hour for three people) within the entire sampling plot. Earthworms of all life stages (adults, juveniles and cocoons) were collected and kept in 95 to 100 % ethanol. Ethanol was changed after 24 hours to ensure clean fixation.

For soil analyses, ten soil cores per plot were collected at 0–10, 10–20 and 20–30 cm depth, each 20 cm along a transect through the sampling plot at Galbao, Itoupé, Mitaraka and Trinité (Vleminckx et al., 2019). Cores were combined into a composite 500 g sample, which was dried at 25 °C and sieved at 2 mm. Physical and chemical analyses were done at CIRAD Laboratory (Montpellier, France, (“Unité de service Cirad - Analyses,” n.d.)) with protocols available on their website (Pansu and Gautheyrou, 2007). Measured variables were soil texture (clay, fine silt, coarse silt, fine sand and coarse sand), pH H_2_O, organic carbon, total nitrogen, C/N ratio and available phosphorus (Supplementary Table 1).

We also collected nine other climatic variables and soil properties from global databases. We used seven climate layers from the CHELSA climate database (Karger et al., 2017) corresponding to temperature and precipitation variables (annual mean temperature, temperature seasonality, temperature annual range, annual precipitation and precipitation seasonality). In addition we averaged the values of the first 30 cm for bulk density and cation exchange capacity (CEC) obtained from the SoilGrids database (Hengl et al., 2017).

### 2.3. DNA barcoding

Earthworms specimens were sorted into morpho-groups (i.e. groups of individuals of similar size, pigmentation and general external morphology) as a conservative approximation of the taxonomic diversity in a sample. We did not attempt to group into the same morphospecies immature stages (i.e. cocoons or juveniles) and adults, because the former usually lack any reliable character to link them with the corresponding adults. Consequently, cocoons, juveniles and adults were systematically assigned to different morphospecies even when obviously belonging to the same species. Subsequently, up to five specimens per morpho-group for each sample, and all the cocoons and fragments, were selected for DNA barcoding. A small piece of cutaneous tissue (about 1 mm^2^), or of the embryo in case of cocoons, was fixed in ethanol (100 %) and stored at −20 °C before DNA extraction.

Lab work followed standardized protocols for DNA extraction, barcode amplification and sequencing (deWaard et al., 2008). DNA was extracted using a glass-fiber column based protocol (Ivanova et al., 2006). The primer cocktail C_LepFolF and C_LepFolR (Hernández Triana et al., 2014) was used to amplify a 658 bp fragment of the COI gene. The PCR thermal regime consisted of an initial denaturation at 94°C for 1 min; five cycles at 94°C for 1 min, 45 °C for 1.5 min and 72 °C for 1.5 min; 35 cycles of 94°C for 1 min, 50 °C for 1.5 min and 72 °C for 1 min followed by a final cycle at 72 °C for 5 min. Each PCR product was cleaned up using Sephadex (Ivanova and Grainger, 2007). PCR amplicons were visualized on a 1.2% agarose gel E-Gel® (Invitrogen) and then diluted 1:10 with sterile water. Amplicons (2–5 μL) were bidirectionally sequenced using sequencing primers M13F or M13R (Messing 1983) and the BigDye® Terminator v.3.1 Cycle Sequencing Kit (Applied Biosystems, Inc.) on an ABI 3730xl capillary sequencer following manufacturer’s instructions. All sequences and supporting information obtained for Galbao, Itoupé, Mitaraka and Trinité were combined with the ones obtained from samples taken at the Nouragues stations during a previous study (Decaëns et al., 2016) and deposited in the Barcode of Life Datasystems (BOLD) database (Ratnasingham and Hebert, 2007) in the dataset “Earthworms from the tropical rainforest of French Guiana” (DS-EWFG, DOI: dx.doi.org/10.5883/DS-EWFG).

### 2.4. Delimitation of the Molecular Operational Taxonomic Units (MOTUs)

MOTUs delimitation was done using the Automatic Barcode Gap Discovery method (ABGD, Puillandre et al., 2012). We did not use the BIN (Barcode Index Number) delimitation system from BOLD (Ratnasingham and Hebert, 2013) because as shown by Decaëns et al. (2016) who compared different methods, this one seems to not be suitable for the case of earthworms which presents a higher divergence rate compare to other groups. In a first step we used only the sequences from Nouragues and an *a priori* threshold of 12 - 14 %, to verify recovery of the 48 MOTUs found in the original study by Decaëns et al. (2016). The ABGD parameters obtained in this first run (p = 0.05, P = 0.2, relative gap width X = 0.5, distance d = K80) were subsequently applied to the full dataset comprising sequences from all locations in order to delimited new MOTUs.

### 2.5. Data analyses

Data from Nouragues locations were only used to delimit the MOTUs and assess the diversity at the regional and local scales. All other following analyses were only performed for Galbao, Itoupé, Mitaraka and Trinité, as a complete dataset including soil properties was available only for these four locations and because only qualitative sampling has been performed at both Nouragues location in 2016. Also, under the category “hilltop forests” we grouped the inselberg-like habitats (i.e. hilltop and transition forests and rocky savannah) because they shared some soil characteristics and for many of this three habitats, the number of replicates was not enough to analyse them separately.

#### 2.5.1 Alpha-to gamma-diversity estimates

To compare species diversity among different localities, habitats and microhabitats, we adjusted rarefaction and extrapolation curves for MOTU diversity using the “iNEXT” package (Hsieh et al., 2016) for R (R Core Team, 2020). At the regional scale, we used the observed and extrapolated number of MOTUs according to the number of sampling locations as a measure of sampling effort; whereas at local scale we used the number of specimens collected to account for the variability in earthworm density among habitats and microhabitats. We further computed observed richness, defined as the number of different MOTUs observed for each locality, habitat or micro-habitat, as well as the estimated species richness (Chao estimate), using the R package “vegan” (Oksanen et al., 2019).

Also, we looked at the diversity captured by each of the three sampling methods and in each kind of microhabitats, and by drawing Venn diagrams using the package “eulerr” (Larsson, 2020). This enabled us to highlight the proportion of MOTUs shared between sampling methods and microhabitats.

#### 2.5.2. Beta-diversity analyses

We calculated the average Sorensen’s index of dissimilarity using the R package “betapart” (Baselga et al., 2018) to assess the variation of MOTU composition among sites (β-diversity) and its decomposition into spatial turnover (i.e. replacement of species from location to location) and nestedness (i.e. when the composition of a site is a subset of another site hosting more species) (Baselga, 2010). This was done at three different levels to comprehensively characterise the relative contribution of distance and habitat diversity to β-diversity: (1) by adopting the same approach for each habitat separately (i.e. *between localities + within habitat* by comparing with each other the four species lists found in each locality for a given habitat); (2) the ecological β-diversity by comparing the composition of different habitat species pools within each locality (i.e. *within locality + between habitats*); (3) the local β-diversity by comparing the composition of individual communities collected in a given habitat at a given locality (i.e. *within locality + within habitat*).

Singletons are MOTUs represented by only one individual in the dataset, and, because they are by definition present only in a single location, they are expected to inflate the indices of β-diversity when present at a high proportion. To account for this potential bias, we performed all analyses with and without singletons. As we could not find any significant differences, we decided to keep singletons present in the analyses presented herein.

#### 2.5.3 Environmental drivers of community composition

Data were organized into two separate tables to perform a transformation based canonical redundancy analysis (tb-RDA, Legendre and Gallagher, 2001), in order to highlight the environmental parameters explaining the observed variations in earthworm community composition. This analysis was done using the *rda* function of the R package “vegan” (Oksanen et al., 2019) for a subset of 32 sampling plots from Galbao, Itoupé, Mitaraka and Trinité, representing replicated habitats (mainly hilltop, plateau, slope and swamp forests) for which soil variables were available (Supplementary Table 1). We used for this abundance data as a contingency community table composed of 32 rows (i.e. the 1 ha sampling plots, see Supplementary Table 1) and 81 columns (i.e. the MOTUs), and another table with the same 32 rows which contained the explicative variables grouped as spatial, soil texture, soil chemistry and climatic variables (see Supplementary Table 2 for detail). MOTU abundance data were Hellinger-transformed before computing the tb-RDA to reduce the weight of the most abundant groups in the analyses. After removing the variables that were correlated based on Pearson correlation, we used the function *ordiR2step* to select the explanatory variables that contribute significantly to the model and permutation tests to verify the significance of the RDA model obtained. Finally, we looked at the relative contribution of the different groups of explicative variables (spatial / soil texture / soil chemistry) using a partial RDA ordination with the function *vpart* of the “vegan” package (Borcard et al., 2018).

## 3. Results

### 3.1. Barcoding results and MOTUs designation

A total of 1819 earthworm specimens out of 55 sampling points from the four localities of Galbao, Itoupé, Mitaraka and Trinité were selected for DNA barcoding. We were able to obtain 1683 COI sequences (after removal of contaminated sequences), with a sequencing success of 92.52 %.

After adding the dataset from the Nouragues (Decaëns et al., 2016) and removing the sequences shorter than 300 bp, our total dataset contained 2304 sequences (409 from Galbao, 595 from Itoupé, 347 from Mitaraka, 324 from Trinité, 431 from Nouragues-Inselberg and 198 from Nouragues-Pararé) clustering into 119 MOTUs when using a 13 % threshold on ABGD (Figure 2). The dataset comprised 821 adults (35.6 %), 1 276 juveniles (55.4 %), 119 cocoons (5.2 %) and 88 fragments (3.8 %). Mean intra-MOTU divergence was 2.12 % (ranging from 0 % to 9.26 %) and mean inter-MOTU was 24.02 % (ranging from 10.56 % to 49.98 %) (Figure 2). We found 30 MOTUs (25.2 % of the total number) solely represented by juveniles, cocoons and specimen fragments. In total, 24 MOTUs (20.2 % of the total number) were singletons, 13 of which were represented only by juveniles and one by specimen fragments.

**Figure 2:**
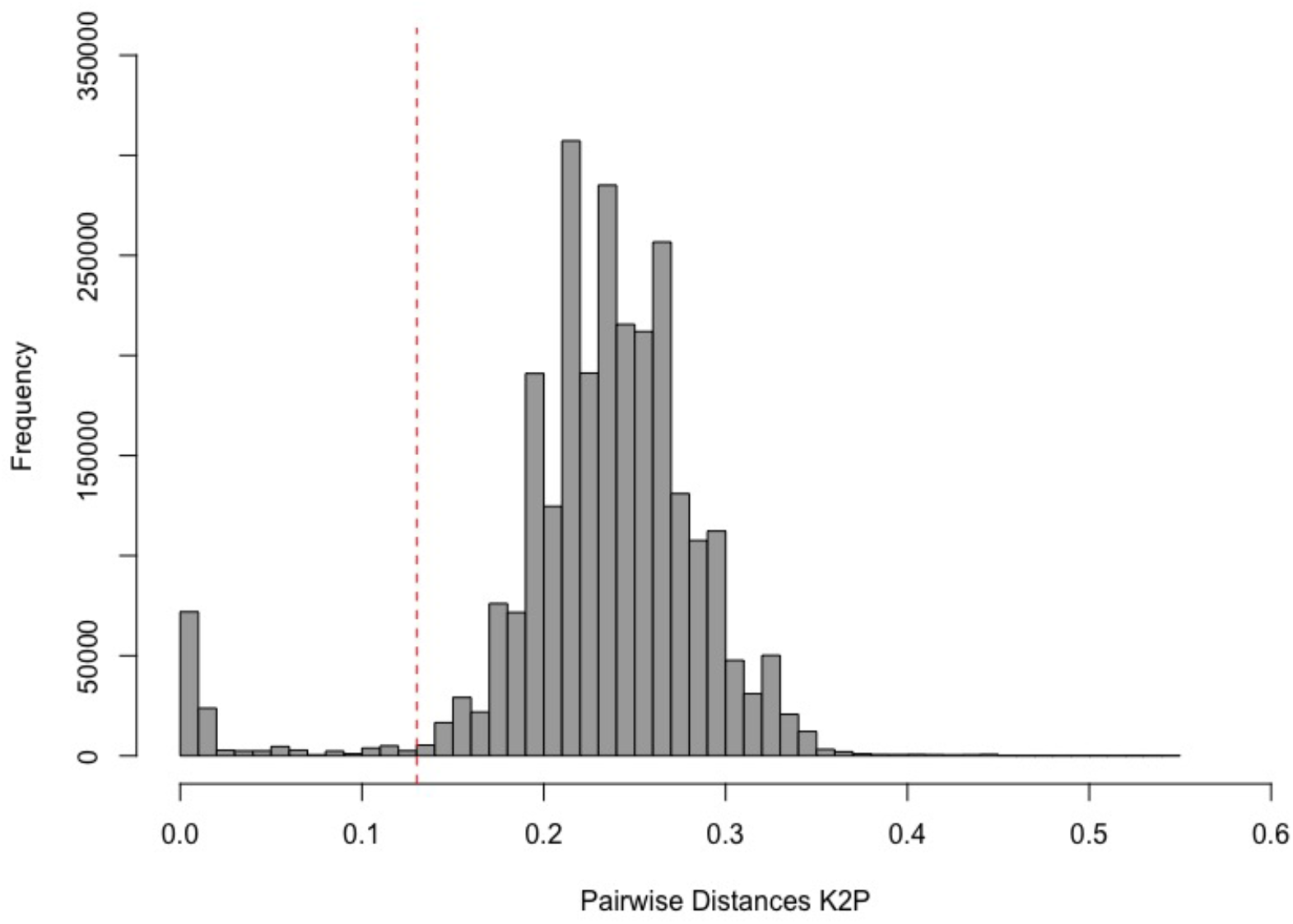
Barplot showing the pairwise distance distribution of 2304 DNA barcode sequences. The dotted red line represents the threshold value between intra-MOTU (left) and inter-MOTU (right) pairs of individuals, used in ABGD (13 %).

### 3.2. Earthworm diversity at regional scale

There were only a few shared MOTUs between the sampled locations; only two (1.7 %) MOTUs were shared between the six locations and 85 (71.4 %) were present at a single location (Figure 3). Localities that shared the most MOTUs also seemed to be closer geographically, such as both Nouragues locations (~ 5.60 km) that shared 18 MOTUs or Nouragues Inselberg and Galbao (~ 87 km) which shared 10 MOTUs (Figure 3B). However, even distant localities such as Trinité and Mitaraka (~ 285 km) can share five MOTUs (Figure 3B). As a consequence, the rarefaction and extrapolation curve fitted for the full dataset at regional scale shown a sharp increase of MOTU counts with increasing numbers of sampling locality (Figure 4). There was no evidence for any saturation of the regional species pool, even when extrapolating the number of MOTUs that would result from doubling the sampling effort, and asymptotic richness estimates suggested that more than 250 species could occur at this scale (Table 1). Our results therefore indicated that the 119 observed MOTUs that we found during our survey might represent only less than half (45.4 %) of the real number of earthworm species that may exist in the entire French Guiana. However, there was a large uncertainty for this estimate (SD = 45.7).

**Figure 3:**
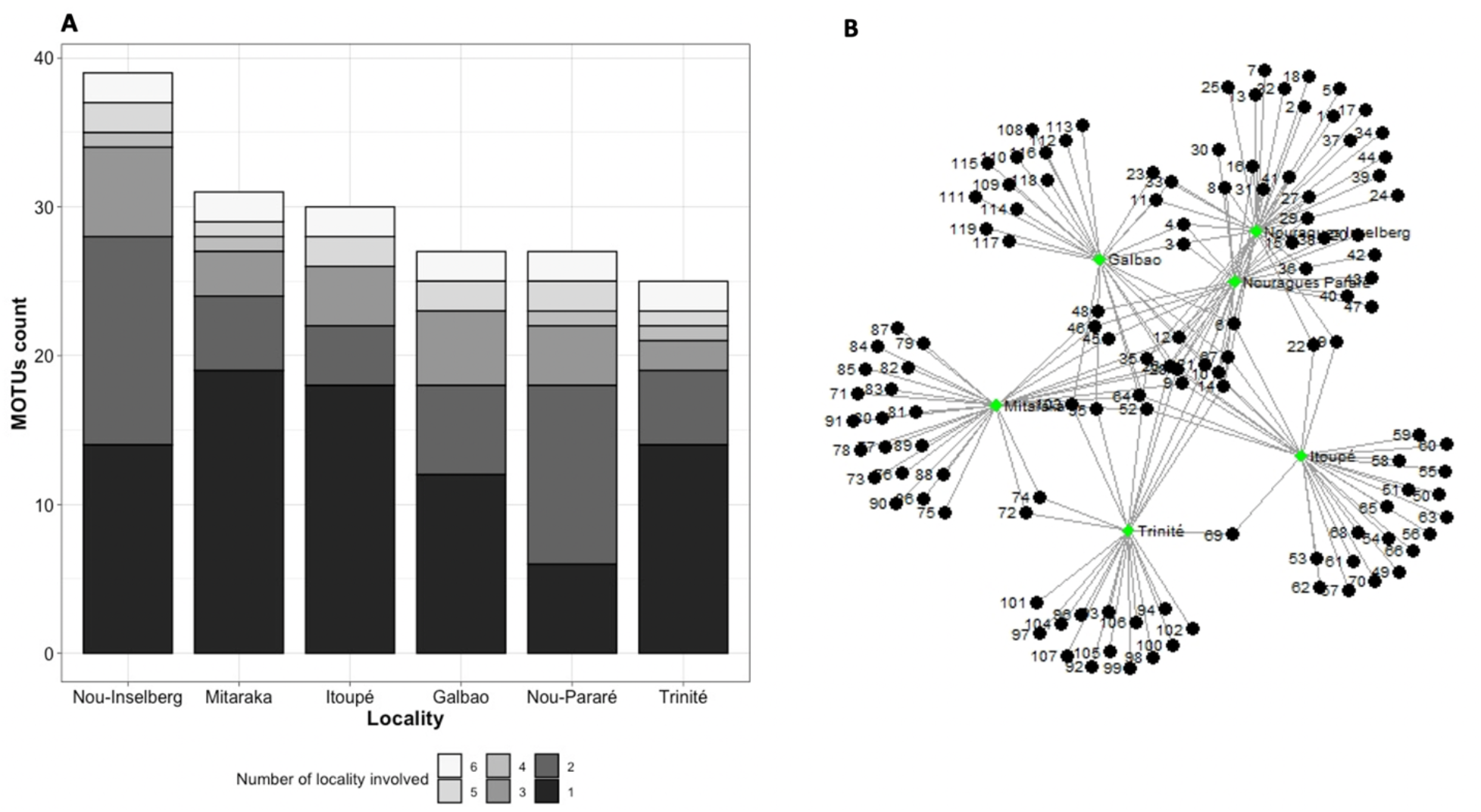
A) Barplot showing the total richness for each locality as well as the part that is specific to one location (1) and shared between 2 to 6 locations. B) Bipartite network of the MOTUs showing detail on how the MOTUs are shared between the four locations. Each circular node represents a MOTU and each square a locality. (Nou = Nouragues).

**Figure 4:**
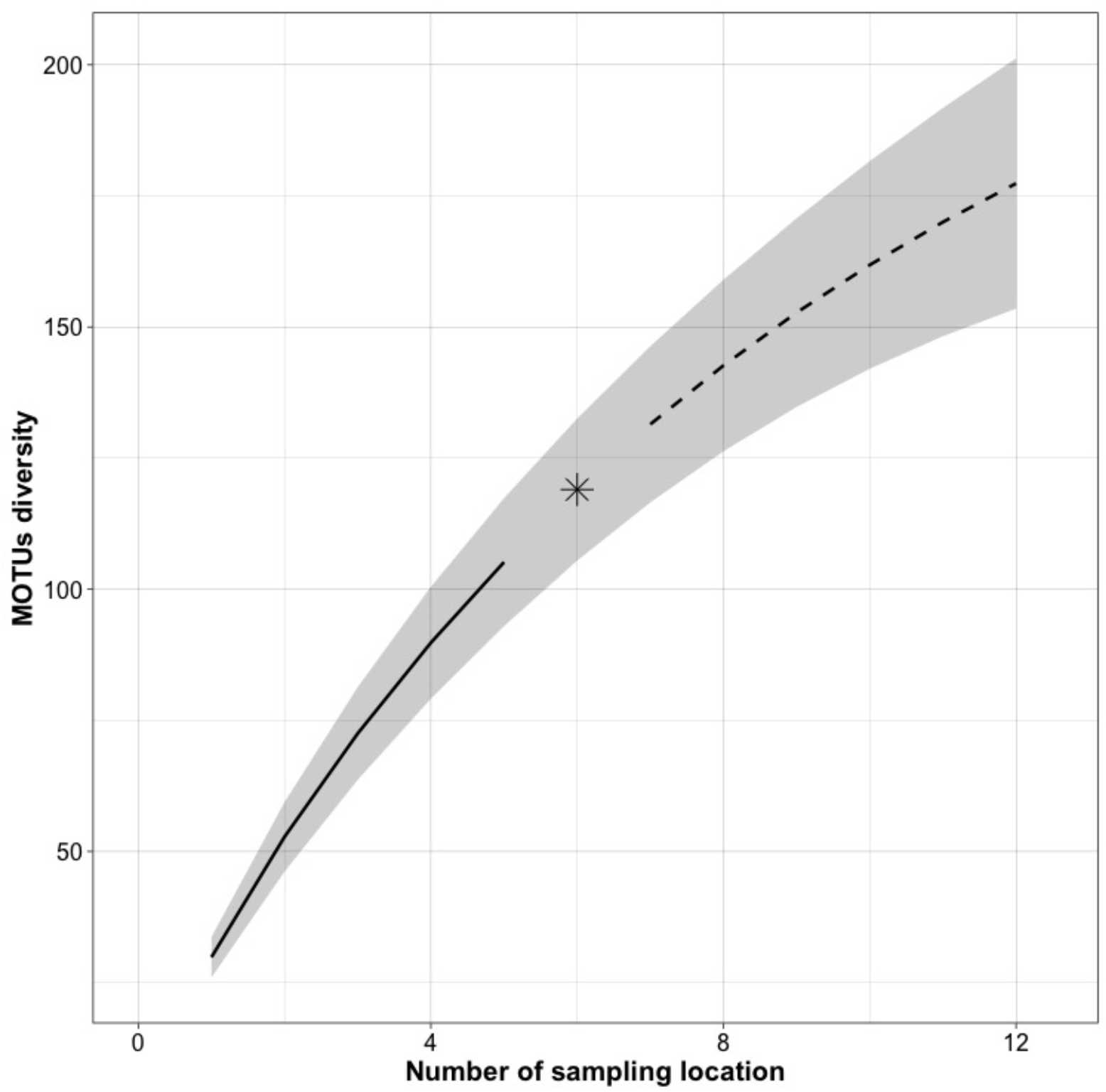
Rarefaction and extrapolation curves showing MOTUs accumulation according to the number of sampled locations. A total of 119 MOTUs were observed for six sampled locations (Galbao, Itoupé, Mitaraka, Trinité and Nouragues-Inselberg and Nouragues-Pararé). Solid line corresponds to rarefaction curve, dashed line to extrapolation curve; shaded area represents a 95% confidence intervals based on a bootstrap method with 200 replications.

**Table 1:** Number of COI sequences per locality and associated diversity indices, calculated with the package *vegan* Indices were calculated using MOTUs as species proxy. The last row “All” represents observed and estimated richness at the regional scale, for all six localities. Abbreviations: S.obs = observed richness, S.chao1 = index of estimated richness and se.chao1 = standard error for s.choa1.

| Localities | # of sequences | S.obs | S.chao1 | se.chao1 |
| --- | --- | --- | --- | --- |
| Galbao | 409 | 27 | 30.5 | 3.65 |
| Itoupé | 595 | 30 | 32 | 2.58 |
| Mitaraka | 347 | 31 | 33.5 | 2.89 |
| Trinité | 324 | 25 | 26.5 | 2.22 |
| Nouragues_Inselberg | 425 | 39 | 50.14 | 8.23 |
| Nouragues_Pararé | 204 | 27 | 36.33 | 8.84 |
| All | 2304 | 119 | 262.35 | 45.70 |

The Sorensen indices of regional beta-diversity (β_SOR_, *between locality + within habitat*) were high and similar when comparing different forest habitats, showing a strong spatial turnover (β_SIM_) at this scale (Supplementary Figure 1).

### 3.3. Earthworm diversity at local scale

Rarefaction and extrapolation curves for individual locations shown how MOTUs accumulate as a function of the number of sampled individual (Figure 5). The observed richness ranged from 25 MOTUs in Trinité to 39 in Nouragues-Inselberg (Table 1). For almost all locations, richness estimates appeared close to the observed values, except for both Nouragues locations where the estimated richness was 1.3 times higher than the observed total richness. The standard error of the Chao index for both Nouragues locations was also higher than for the four other locations. These observations reflected the trend that the slope of the rarefaction curves of Nouragues Inselberg and Pararé seemed more pronounced than that of the slopes from other localities. However, none of the localities seemed to reach an asymptote.

**Figure 5:**
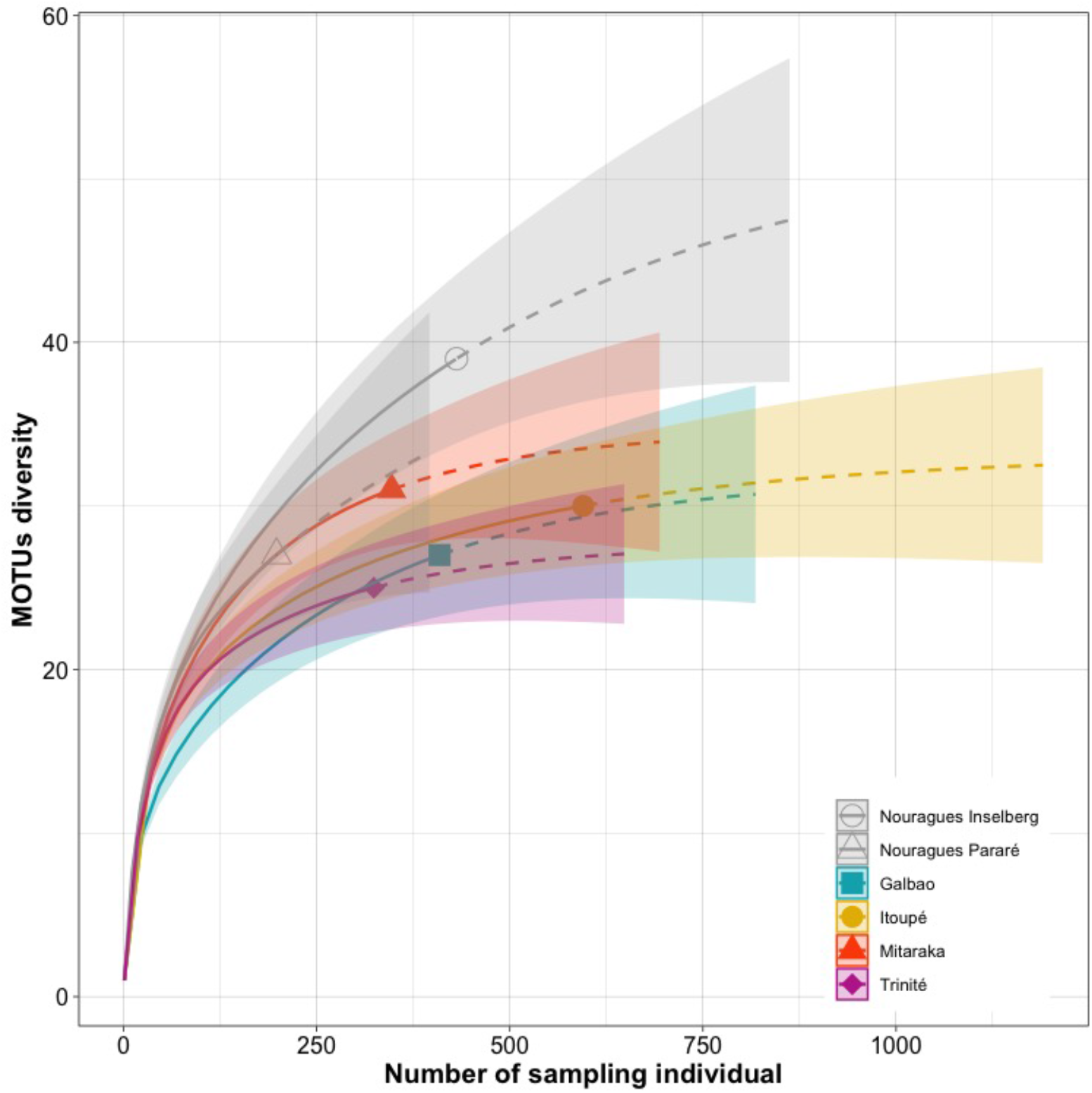
Rarefaction and extrapolation curves showing MOTU accumulation according to the number of sampled individuals in the six different locations separately. Rarefaction curves are represented in solid lines, extrapolation curves in dashed lines; shaded areas represent a 95% confidence intervals based on a bootstrap method with 200 replications.

**Figure 6:**
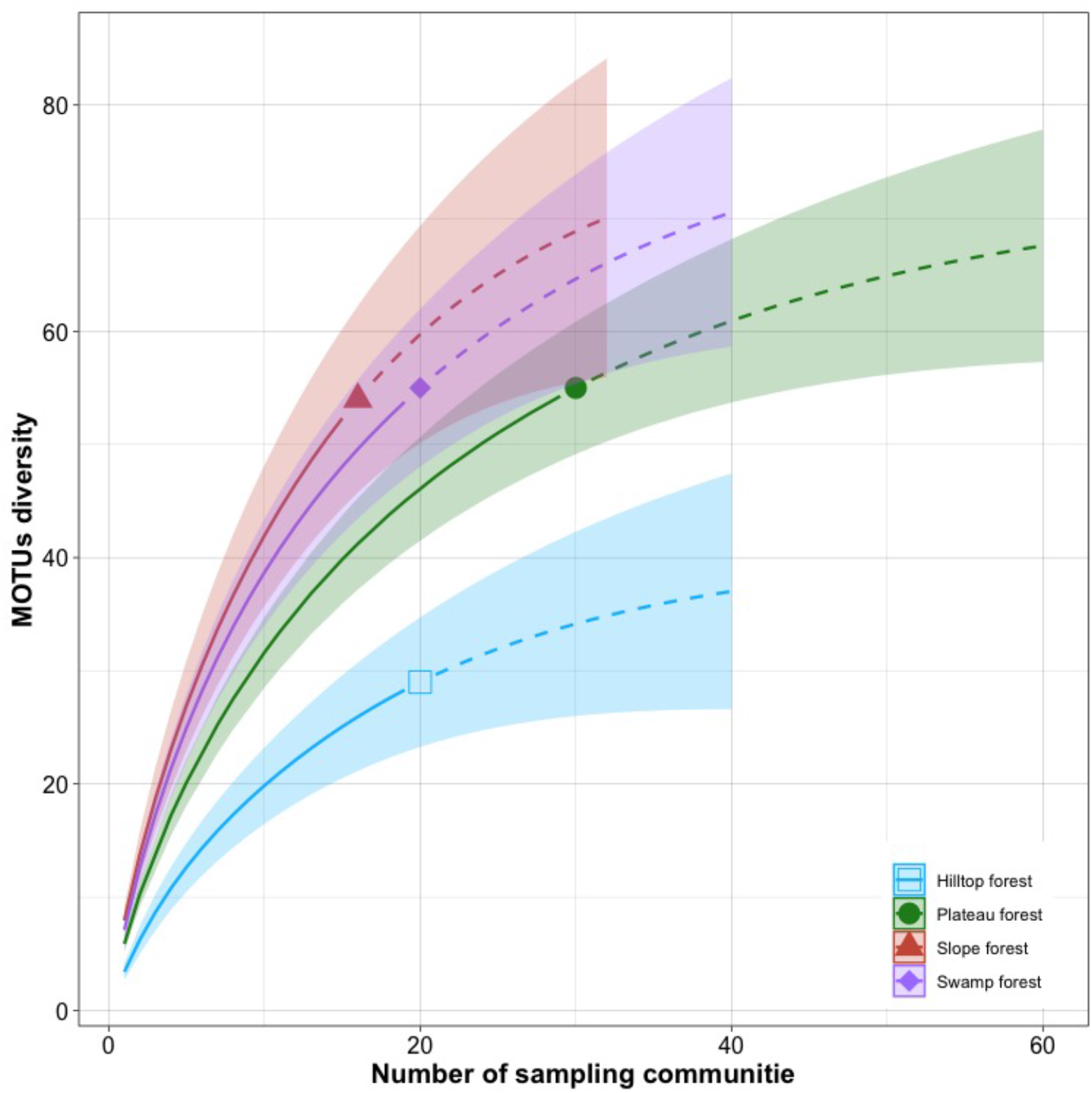
Rarefaction and extrapolation curves showing the MOTU accumulation according to the number of sampled communities for each main type of habitat at the regional scale, from all six available localities. Rarefaction curves are represented in solid lines, extrapolation curves in dashed lines; shaded areas represent a 95% confidence intervals based on a bootstrap method with 200 replications.

The Sorensen indices of beta-diversity (β_SOR_) calculated at local scale among the different habitats inside each of the four new locations (*within habitat + within locality*) were fairly variable (Supplementary Figure 2), but all showed a partitioning in favour of spatial turnover (β_SIM_) in comparison to nestedness (β_SNE_). Overall, the ecological beta-diversity (β_SOR_ *within locality + between habitat*) among different habitats was high and similar when comparing the different localities, ranging from 0.62 from Galbao to 0.69 in Mitaraka (Supplementary Figure 3). However, it was explained in greater part by nestedness in Itoupé (0.44), and by turnover in Galbao, Mitaraka and Trinité (0.55, 0.51 and 0.44 respectively).

### 3.4. Earthworm diversity at habitat level

Several habitats were sampled with different levels of richness. Overall, at the regional scale, plateau, slope and swamp forests seemed to harbor a similar diversity (with respectively 55, 54 and 55 MOTUs) higher than the diversity in hilltop forest (29 MOTUs). The extrapolation did not show sign of saturation at the regional scale. However, these observed trends were not necessarily conserved at the local scale, where the differences in diversity between habitats could be more pronounced and for certain localities the rarefaction curves for some habitats seem to reach a plateau. Indeed. a higher richness was observed for the hilltop forest in Galbao, the slope forest in Itoupé, the plateau forest in Mitaraka and the swamp forest in Trinité and both Nouragues localities (Supplementary Figure 4). However, for Itoupé, Trinité and Nouragues Pararé, this was confounded by the number of individuals sampled. The rarefaction curves of some habitats, such as the plateau forest in Galbao and Nouragues Inselberg and the slope forest in Itoupé, almost reached an asymptote with a narrow standard error (Supplementary Figure 4). However, at this scale, a substantial part of the MOTUs are not shared between habitats of the same location.

### 3.5. Earthworms diversity at microhabitat level and sampling comparison

On average, between 11.2 and 46.5 earthworms per meter square (SD: 28.4 – 54.6) were collected per sampling plot with the TSBF method only depending on the locality, with Mitaraka showing the lowest and Trinité the highest abundance (Supplementary Figure 5A). The richness of MOTUs recovered with this method (TSBF) was also quite variable, as it ranged on average from 0.3 in Mitaraka to 1.3 on Galbao (SD: 0.6- 1.1), with a regional mean of 0.9 (SD = 1.2) per sampling plots (Supplementary Figure 5B). However, when using all sampling methods combined, we found a higher average of richness (Supplementary Figure 5B) at local scale. The regional mean richness was 6.1 (SD = 3.1) all sampling methods combined (Supplementary Figure 5B). This trend is also observable at the habitat scale (Supplementary Figure 5 C & D).

Overall, the qualitative sampling approach allowed the collecting of roughly twice the number of MOTUs that was recovered by the quantitative TSBF method alone (Figure 7A). All but six of the 90 MOTUs found at Galbao, Itoupé, Mitaraka and Trinité were collected by qualitative sampling, while the quantitative sampling only resulted in the finding of 42 MOTUs (Figure 7A). Most of the diversity recovered by all sampling methods was in the soil with 69 MOTUs, but 52 MOTUs were however found in other types of microhabitats (Figure 7A). Also, among the 119 MOTUs found with the full dataset, 23 were exclusively found in other microhabitats than soil (mostly decaying trunk) (Figure 7B).

**Figure 7:**
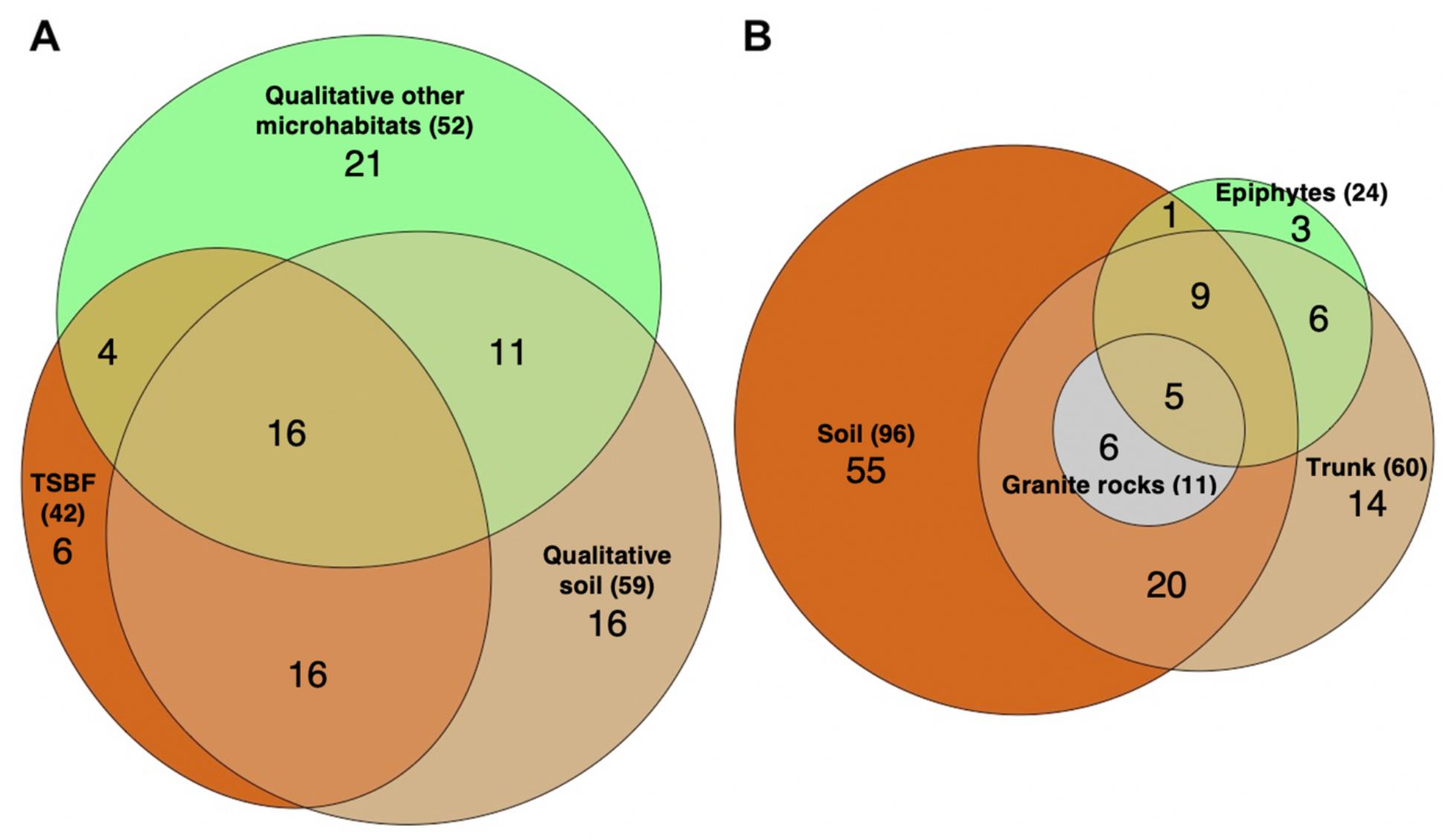
Venn diagram showing the number of MOTUs recovered by A) the different sampling methods (TSBF = quantitative) performed in Galbao, Itoupé Mitaraka and Trinité only (total MOTU = 90), as only the qualitative method has been used in the Nouragues; and B) the different microhabitat types sampled with the whole range of sampling methods and all the dataset (total MOTU = 119). Numbers in brackets are the total number of MOTUs (i.e. unique and shared) found for a given sampling method or microhabitat.

### 3.6. Community composition

The climatic variables, as well as bulk density and cation exchange capacity, were not retained in the RDA analysis because they resulted to be highly correlated with the spatial variables (i.e. elevation and precipitation). The *ordiR2step* function selected all spatial variables (topography, longitude, latitude and elevation), fine silt content, pH, total nitrogen, organic carbon and phosphorus as subset of explanatory variables to explain earthworm community composition (Supplementary Table 2).

Communities in Galbao, Itoupé, Mitaraka and Trinité were generally well separated by the first four RDA axes, as well as the forest types within each location (Figures 8). With the exception of swamp forests from Galbao, Mitaraka and Trinité and plateau and hilltop forests from Galbao and Trinité that could be observed close together on the ordination space, forest types between localities rarely seemed to share similarities in their composition and environmental properties. The first two and first four canonical axes together explained 22.6 % and 36.4 % respectively of the total variance of the response data, with an adjusted R^2^ of 0.3697. The variable elevation played an important role in the distribution of the sampling plots along the first axis (Figure 8A). Higher elevation values were observed at Galbao and Itoupé, and the lowest at Trinité. The variables fine silt content and hilltop topography were correlated with the second axis (Figure 8A). Soils at Galbao contained high silt content. The pH was correlated with the third axis. And, the fourth axis opposed the latitude and longitude to organic carbon, total nitrogen and phosphorus (Figure 8C and 8E). Soils at Mitaraka showed highest content of chemical variables and Galbao the lowest.

**Figure 8:**
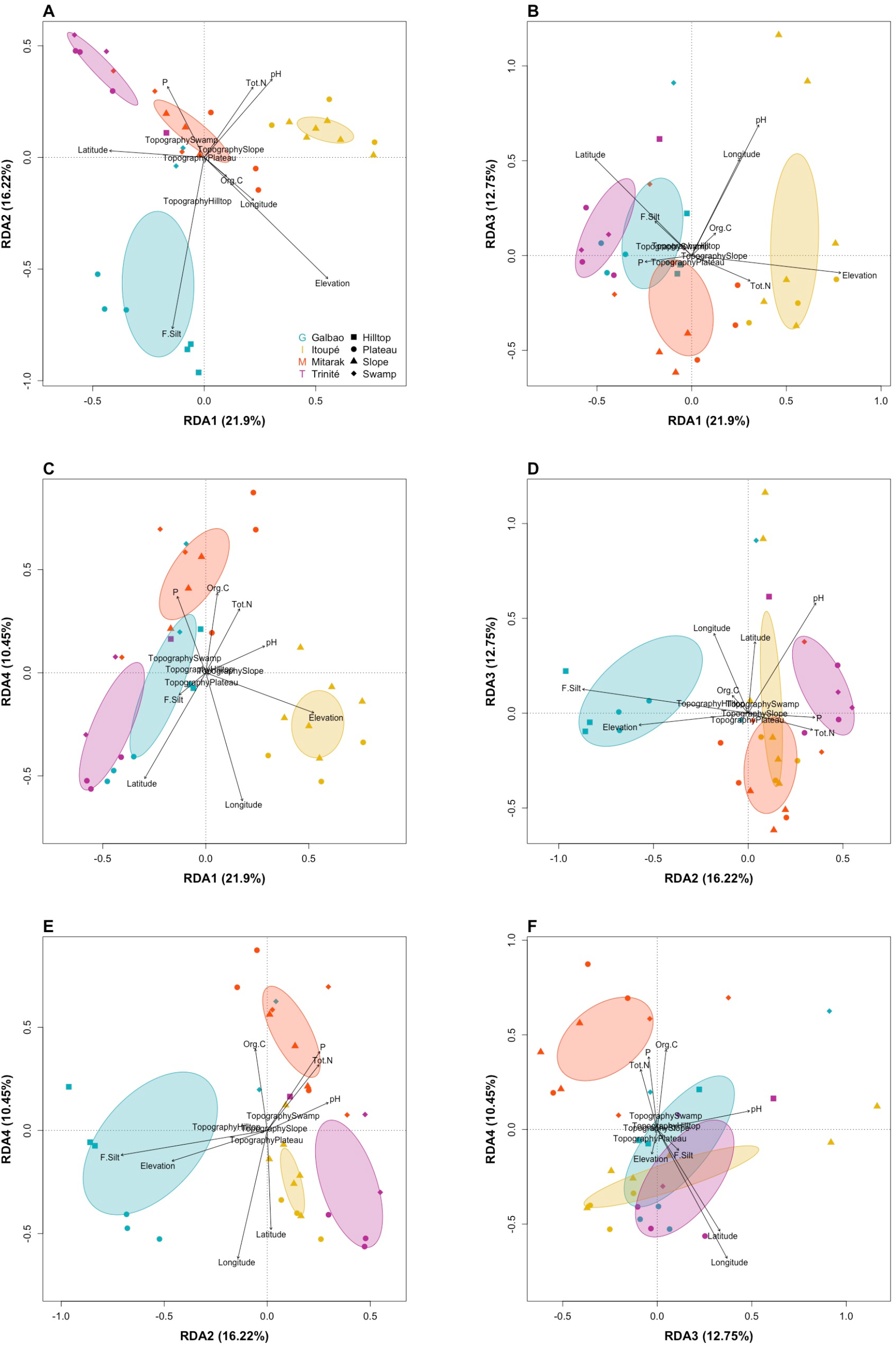
MOTU-environment triplot of RDA after variables selection, showing the relationship between MOTUs composition and environmental data (scaling 1) on the first two axes (A); the relationship between the communities and the environmental data (scaling 2) on the first two axes (B); the axis 1 and 4 (C); and the axis 2 and 4 (D). Arrows represent the quantitative environmental variables and their length indicates the correlation between the environmental variables and the ordination axes. Abbreviations: C.Sand = coarse sand, C.Silt = coarse silt, P = available phosphorus and Topo = Topography.

The partial RDA showed that spatial variables (elevation, longitude, latitude and topography) explained 19.2 % of the variance in the earthworm community composition, while soil chemical variables (pH, organic carbon, total nitrogen and phosphorus) and soil texture (fine silt content) accounted for 10.8 % and 6 % respectively. The remaining 63 % corresponded to the residuals, i.e. the fraction of the overall variance that was not explained by our selected environmental variables. This meant that the model might still displayed some dominant residual structure.

## 4. Discussion

### 4.1. Unmatched levels of earthworm diversity in French Guiana

To date, only a few publications have addressed the diversity of earthworms at regional scale in French Guiana and the neotropical region overall. In 2007 Brown and Fragoso listed 33 species on the French Guiana territory, but 12 of them have not been described. In 2012, Pavlicek & Csuzdi added one new species to the 21 previously described species from Brown and Fragoso (2007), while suggesting that this figure probably reflected nothing else than the extent of the lack of knowledge on the subject. In 2016, Decaëns et al. published a study based on an approach similar to ours, in which they described the local distribution of 48 MOTUs in the Nouragues reserve. Our results represent a significant step forward in the acquisition of knowledge on this subject, as we were able to detect 119 MOTUs in six study locations distributed all over the region. Furthermore, the rarefaction curve and richness estimates computed at the regional scale both indicate we still only recovered ~ 45 % of the true regional diversity, and that any newly added sampling location could lead to the addition of about 15 to 30 new MOTUs.

The mean observed diversity of 32.2 MOTUs (SD = 9.1), potential species, per location is one of the highest reported for earthworm communities in tropical forests. For comparison, 10 to 17 species have been reported at a similar spatial scale in the native forests of Santa Catarina and Rio Grande do Sul, Brazil (Bartz et al., 2014; Steffen et al., 2018), and in the Mexican tropical rainforest of Chiapas (Fragoso and Lavelle, 1987) using s traditional taxonomy approach. We also calculated a mean of 2.7 species (SD = 3.7) per sampling plots with the TSBF sampling method which seems to at least double what would be expected in this region according to the model used in Phillips et al. (2019) (about 1 species in French Guiana / Northern Amazonia).

This discrepancy could be related to the adding value of the sampling methodology we adopted for our earthworm surveys. In earthworm studies, only adult specimens are typically used for species richness assessment, mostly because of the difficulty to identify juveniles to species level. Contrary to this classical approach, we included in our study juveniles, cocoons and fragments of earthworms that represented the majority (64 %) of our dataset. Without them, we would have work with about 36 % of our dataset and would have missed a quarter of the regional diversity (i.e. 30 MOTUs) represented by MOTUs only present in the samples as immatures or fragmentary remains. This stresses the importance of an integrative approach to species richness assessment that includes sampling of all life stages and the use of a molecular identification method such as DNA barcoding (Decaëns et al., 2016; Richard et al., 2010).

An additional explanation for our high diversity compared to the model from Phillips et al. (2019) is that previously published studies generally used soil hand sorting (TSBF) as the only quantitative sampling method, providing only a partial picture of the composition of earthworm communities. We, on the other hand, coupled three sampling methods allowing us to prospect not only the soil, but also other types of microhabitats, thereby increasing the number of species that we were able to detect locally. Indeed, our approach allowed us to collect twice as many species as if we had only used quantitative sampling.

Even more interesting is the discovery of 23 MOTUs that were sampled exclusively in microhabitats other than soil (decaying trunks and epiphytes). Arboriculous earthworm species have been observed in tropical regions (Fragoso and Rojas-Fernández, 1996; Lavelle, 1978; Lavelle and Kohlmann, 1984; Rodriguez et al., 2007), and Decaëns et al. (2016) already documented that as much as 35 % of the total number of species observed in both Nouragues locations may occur at least occasionally in epiphytic soils. In oligotrophic soils of the neotropics, earthworm communities are often dominated by pigmented earthworms that prefer microhabitats where organic matter is concentrated including decaying trunks (Fragoso and Lavelle, 1992; Decaëns et al., 2016). This suggests that a significant part of the earthworm diversity in tropical regions with oligotrophic soils could live in aboveground habitats. Overall, the use of this sampling scheme appeared to be very efficient in discovering and describing earthworms richness in tropical region, as it has been previously highlighted in the context of a study of earthworm richness in agroecosystems in Southern Brazil (Bartz et al., 2014).

### 4.2. Few earthworm species with large geographical ranges

Only two MOTUs (#26 and #28) were present in the six sampled locations, showing a very low level of shared diversity at regional spatial scale. These two MOTUs were also the most abundant species represented by 368 (16 %) and 215 (9.3 %) individuals respectively. MOTU #26 has been identified as *Pontoscolex corethrurus* (Muller, 1856), a peregrine endogeic species that originated from the Guyana shield (a geological formation in northeastern South America that extend over Guyana, Suriname, French Guiana, southern Venezuela, as well as parts of Colombia and Brazil), and which is known to be invasive in a number of other tropical countries (Dupont et al., 2012; Marichal et al., 2010; Taheri et al., 2020). Therefore, it was not surprising that this species has a large range in its own origin area. MOTU #28 has been identified as *Nouraguesia parare* (Csuzdi and Pavlíček, 2011), a large epigeic species supposedly endemic from French Guiana and that is mostly (54 % of the times in this study) found in rotten trunks (Decaëns et al., 2016). These two species have an opposite nature, one being invasive and the other one endemic, but both in their original range show a large spatial distribution. This could certainly be explained by a high dispersal capacity of these two species, allowing them to colonise new habitats more efficiently than others. This has been already described for *P. corethrurus*, which is known to disperse passively with human activities, making it a formidable colonizer regardless of the distance to be traveled (Dupont et al., 2012). In the case of *N. parare*, it is likely that its large size and surface-dwelling behaviour also allows it to move actively over long distances.

Two species from the genus *Wegeneriona*, which is native of South America, were present in the five locations of Galbao, Itoupé, Nouragues Inselberg, Nouragues Pararé plus Mitaraka (MOTUs #12) or Trinité (MOTUs #21). *Dichogaster andina* (Cognetti de Martiis, 1904) (MOTUs #35) was present in the four locations of Nouragues Inselberg and Pararé with Trinité and Mitaraka. This is another invasive species, originating from Africa (Csuzdi et al., 2008). Invasive species are not rare in French Guiana (Lavelle and Lapied, 2003) and even if its presence was already surprising in the Nouragues locations (Decaëns et al., 2016), finding it at locations easily and frequently visited by human could be explained by recent introduction events. However, its presence in a region as remote as Mitaraka was unexpected. This can only be explained by older traces of human activity such as trade between the Amerindian populations who inhabited this area for example, or when the Wayana Indians, who currently inhabit the lower half of the Maroni River and are native to the Caribbean Sea, moved upstream the river a few centuries ago, pushing local tribes towards the Mitaraka Mountains (Fleury et al., 2016).

### 4.3. Outstanding levels of geographical turnover among earthworm communities

About 1/5 of all MOTUs (20.2 %) were singletons, only two MOTUs were shared between all six locations and 83 MOTUs (69.8 %) were present in only one specific location. 34 MOTUs (28.57 %) were shared between two to five locations, sometimes between geographically distant localities and not the direct close locality. This could indicate both a high level of endemism at regional scale with the presence of a significant amount of rare species, and/or a signal of undersampling. Earthworms are known to exhibit higher rates of endemism compared to some other invertebrates groups composing Amazonian biodiversity (Lavelle and Lapied, 2003). French Guiana in particular is characterized by a high coverage of primary forest and a large water network including 840 rivers stretching over a total distance of 112,000 km (“L’office de l’Eau de Guyane,” n.d.). These rivers are often large enough to easily become geographic barriers to earthworm dispersion, leading to the formation of isolated populations and increasing the likelihood of local radiation events as it has been also shown for other taxa (Boubli et al., 2015; Bruschi et al., 2019; Siqueira et al., 2013). This could therefore explain the different species pools that we observed in each study location, and the importance of the spatial turnover component of regional beta diversity.

At local scale, we also found significant levels of spatial turnover, but these were quite variable among habitat. However, they all show that spatial turnover due to MOTU replacement, rather than nestedness, is responsible for this local beta-diversity. Earthworm diversity is known to be at least partly driven by environmental heterogeneity, as previously shown in Mexico (Fragoso and Lavelle, 1987). When looking at ecological scale, our Sorensen indices between locations are very similar indicating comparable variation of composition of different habitat species pools, with Mitaraka showing the highest one. In contrast, the beta-diversity in Itoupé was mostly explained by nestedness and not turnover. Mitaraka harbor large swamp forests that may represent adverse habitats for most of the species occurring in non-flooded forests. Conversely, sampling at Itoupé took place on different altitudinal levels of a single slope of the tabular mountain, with sampling plots relatively close to each other, and not separated by contrasted habitats or large water barriers, when compared to plots in the other locations. These characteristics of these sampling locations could explain the differences observed between them and why we observed a higher nestedness in the ecological beta-diversity at Itoupé.

### 4.4. Spatial and environmental drivers of earthworm community composition

The spatial effect on species composition was also confirmed by the site ordination in our RDA, where each location was represented by a well-resolved cluster as a consequence of spatial turnover. Habitat replicates inside locations were also quite well clustered depending on the axis, showing that there were also environmental influence happening at this scale. However, even if some type of forests seems to share some similarities, we did not observe a regional pattern as same forest types were grouped inside a location but not between locations. The variation partitioning showed that the spatial variables (longitude, latitude, elevation and topography) explained 19.2 % of the variation in species composition which support the importance of spatial turnover being the most important driver. Then the pH, silt content, as well as organic carbon and total nitrogen also significantly contributed at explaining the species composition of earthworm communities. As a result, the spatial variables must play an important role at the regional scale in the variation of the earthworm community composition as shown by the RDA and beta-diversity. And, the environmental variables must play an important role at a lower scale with gradients of organic carbon and silt content as it has been shown before (Fragoso and Lavelle, 1992) and as some of our RDA results suggest, but more sampling points are needed at local scale. It has also been previously shown that temperature and precipitation determine the structure (species richness and abundance) of earthworm communities (Fragoso and Lavelle, 1992; Phillips et al., 2019). While this might be true at global scale, these factors are perhaps less relevant for environments such as tropical rainforests where temperature remains quite constant and where annual precipitation is in the range of 2,000 - 4,000 mm. Here, precipitation and temperature variables were strongly correlated with the elevation and longitude / latitude, so it was hard to separate the effect of each. All our results strongly agreed on the effect of spatial variables, however, increasing the number of sampling plot at local scale would help to investigate the role of environmental and climatic variables in structuring earthworm communities.

Our partial RDA reveled an important part of residual effect. We already mentioned the potential effect of hydrographic network with mechanisms of local radiation in evolutionary history of earthworm communities, generating high spatial turnover in communities’ composition, that could be part of these residual effect. Some other factors not taken into account in this study could also be considered to explain the composition of earthworm communities at different scales. Such as, the surface vegetation, and in particular the litter traits that are known to have a strong explanatory power on the composition of the soil fauna communities in general (Korboulewsky et al., 2016). But also interaction with other soil taxa such as bacteria and fungi could also play a role, and these variables could reduce the residual effect.

## 5. Conclusion

We found a remarkable high regional species richness of earthworms in French Guiana with a high proportion of rare and endemic species and a relatively low (from what would be expected in tropical region compare to temperate region) local species richness. These strong levels of beta-diversity seem to support the remarkable regional diversity observed in French Guiana. And, at the same time, even if our estimates did not converge with those exposed by Phillips et al. (2019), mainly explained by our improved sampling approach, we agreed to conclude that the explosion of species diversity in equatorial ecosystems is verified at the regional scale but not at local scale. Our study confirms the already mentioned usefulness of DNA barcoding to assess the diversity of understudied invertebrates such as earthworms, especially in areas harbouring high diversity such as the tropics. This method allows a fast investigation of the diversity at local and regional level with MOTU clusters as useful species proxy, especially in the absence of further taxonomic information and when investigating earlier life stages and fragmentary remains. However, as mention earlier in the introduction, the use of barcoding is still limited in the study of tropical earthworm communities, resulting in a lack of molecular taxonomic data of tropical earthworm species. Also, the use of a single mitochondrial fragment is not sufficient and more complete data at the genomic level are needed to refine the identification of molecular species. Our study also highlights the importance of sampling design. The inclusion of three sampling methods including the investigation of non-soil microhabitats greatly increased the assessed diversity. On the basis of our results we expect that each additional sample location with different types of habitat would detect 20 to 30 new MOTUs. These results are also the premises showing the richness of earthworm communities in the netropical region.

Further analyses could be performed to investigate the historical and environmental processes leading to the observed spatial patterns. Our results highlight a strong spatial turnover mechanism in the Amazonian region, and the use of functional trait and/or phylogenetic approaches could bring more insights on the assembling rules of earthworm communities in Amazonia. Indeed, it will be interesting to see if the predominant spatial turnover responsible for the variation in the taxonomic beta-diversity described here and in the previous study would also be responsible for the functional and/or phylogenetic beta-diversity.

## Acknowledgements

We are grateful to the Parc Amazonien de Guyane (http://www.parc-amazonien-guyane.fr) and the Réserve Naturelle de la Trinité (http://www.reserve-trinite.fr/) for authorizations and access to the sampling sites. This work was supported by ‘Investissement d’Avenir’ grants managed by the Agence National de la Recherche (CEBA: ANR-10-LABX-25-01; TULIP: ANR-10-LABX-41). In the Mitaraka region, material was collected as part of “Our Planet Reviewed” Guyane-2015 expedition, which was organized by the Muséum national d’Histoire naturelle (MNHN, Paris) and Pro-Natura international in collaboration with the Parc Amazonien de Guyane, with financial support of the European Regional Development Fund (ERDF), the Conseil régional de Guyane, the Conseil général de Guyane, the Direction de l’Environnement, de l’Aménagement et du Logement and the Ministère de l’Éducation nationale, de l’Enseignement supérieur et de la Recherche. Additional funding was provided by the Trinité Natural Reserve. At the Nouragues, the project was supported by two grants CNRS Nouragues’ 2010 and 2011. This study was supported by funding through the Canada First Research Excellence Fund. The funders had no role in study design, data collection, and analysis, decision to publish, or preparation of the manuscript. This work represents a contribution to the University of Guelph Food From Thought research program.

