## Supplementary figures and tables for "At each site its diversity: DNA barcoding reveals remarkable earthworm diversity in neotropical rainforests of French Guiana"

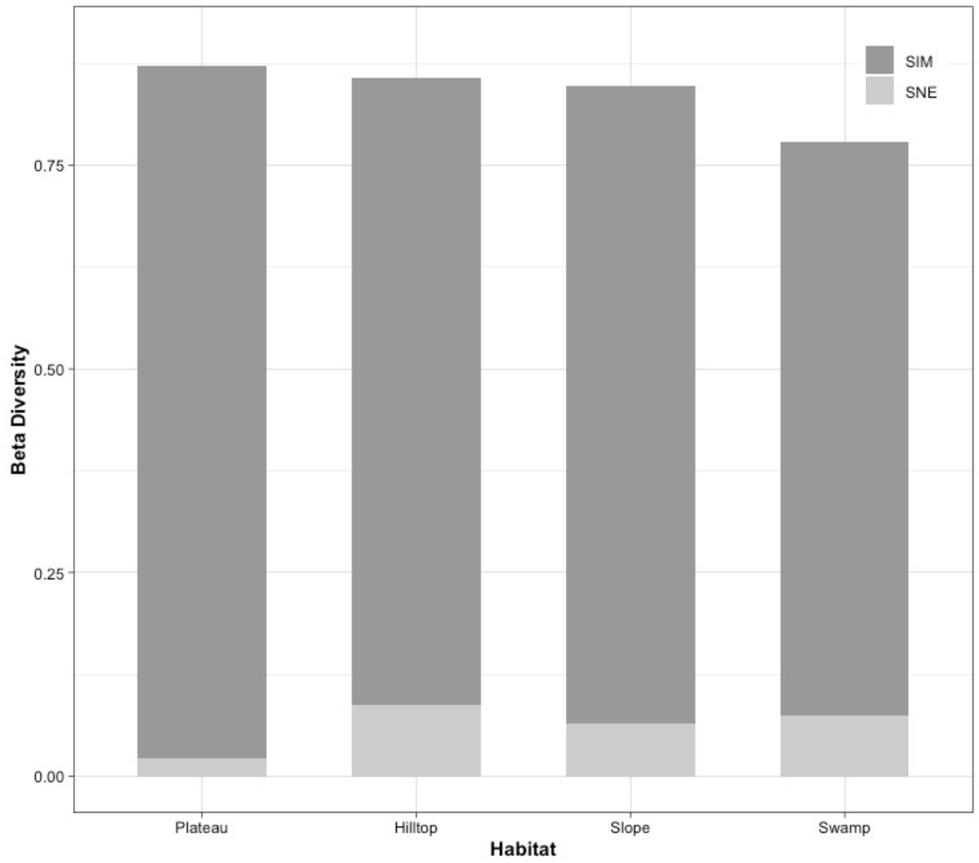


**Supplementary Figure 1:** Partitioning of beta-diversity at regional scale (*between locality* + *within habitat*) into turnover (SIM) and nestedness (SNE).

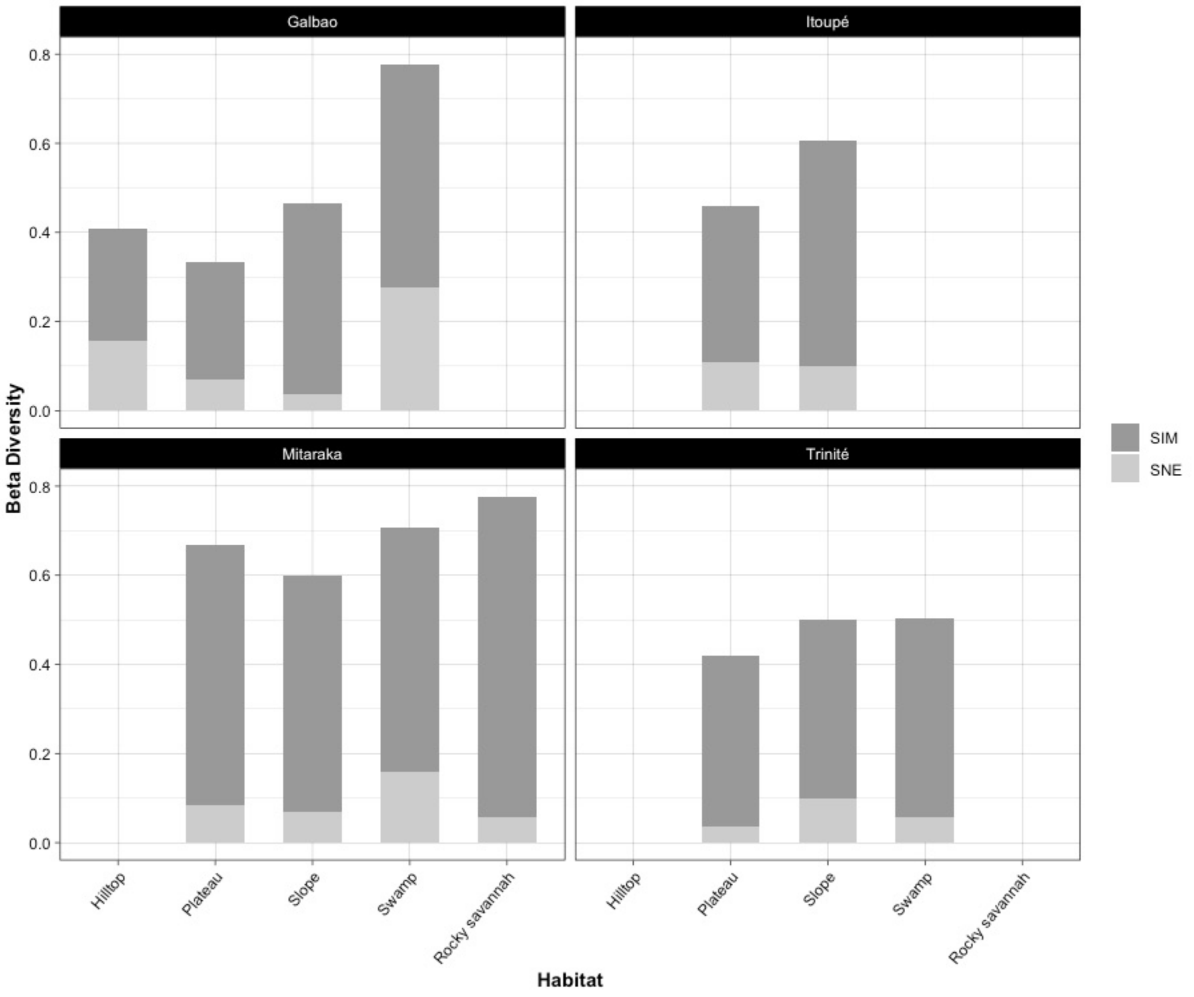


**Supplementary Figure 2:** Partitioning of beta-diversity at local scale (*within locality* + *within habitat*) into turnover (SIM) and nestedness (SNE).

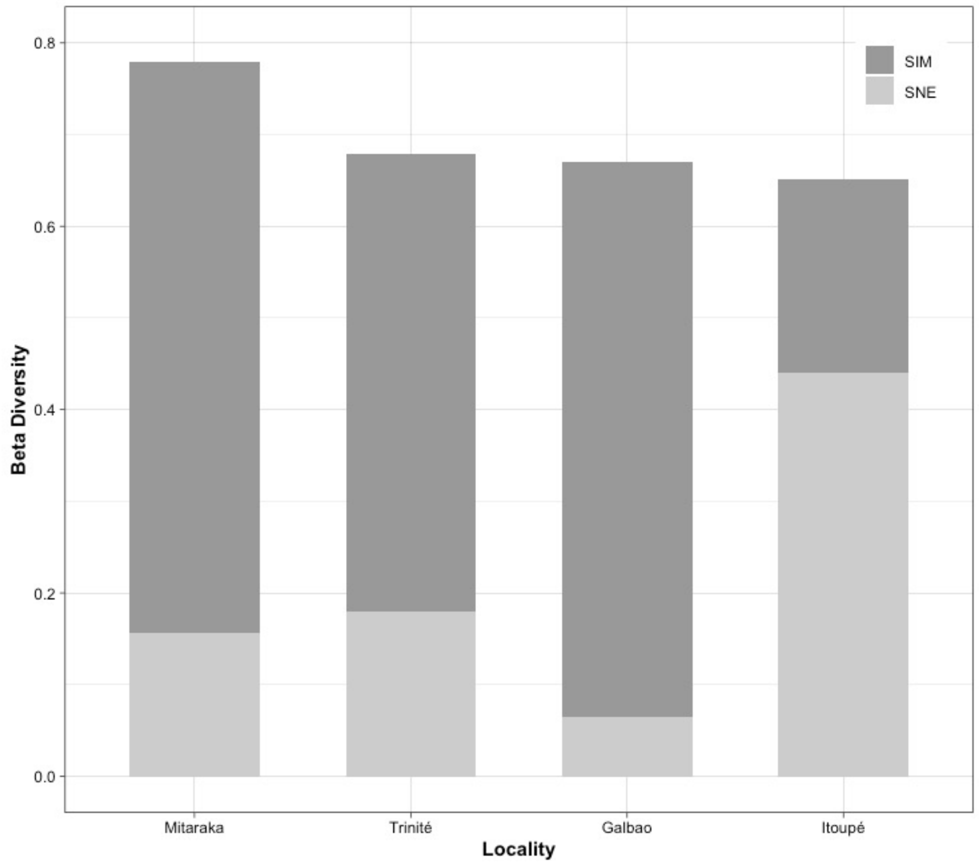


**Supplementary Figure 3:** Partitioning of beta-diversity at ecological scale (*within locality* + *between habitat*) into turnover (SIM) and nestedness (SNE).

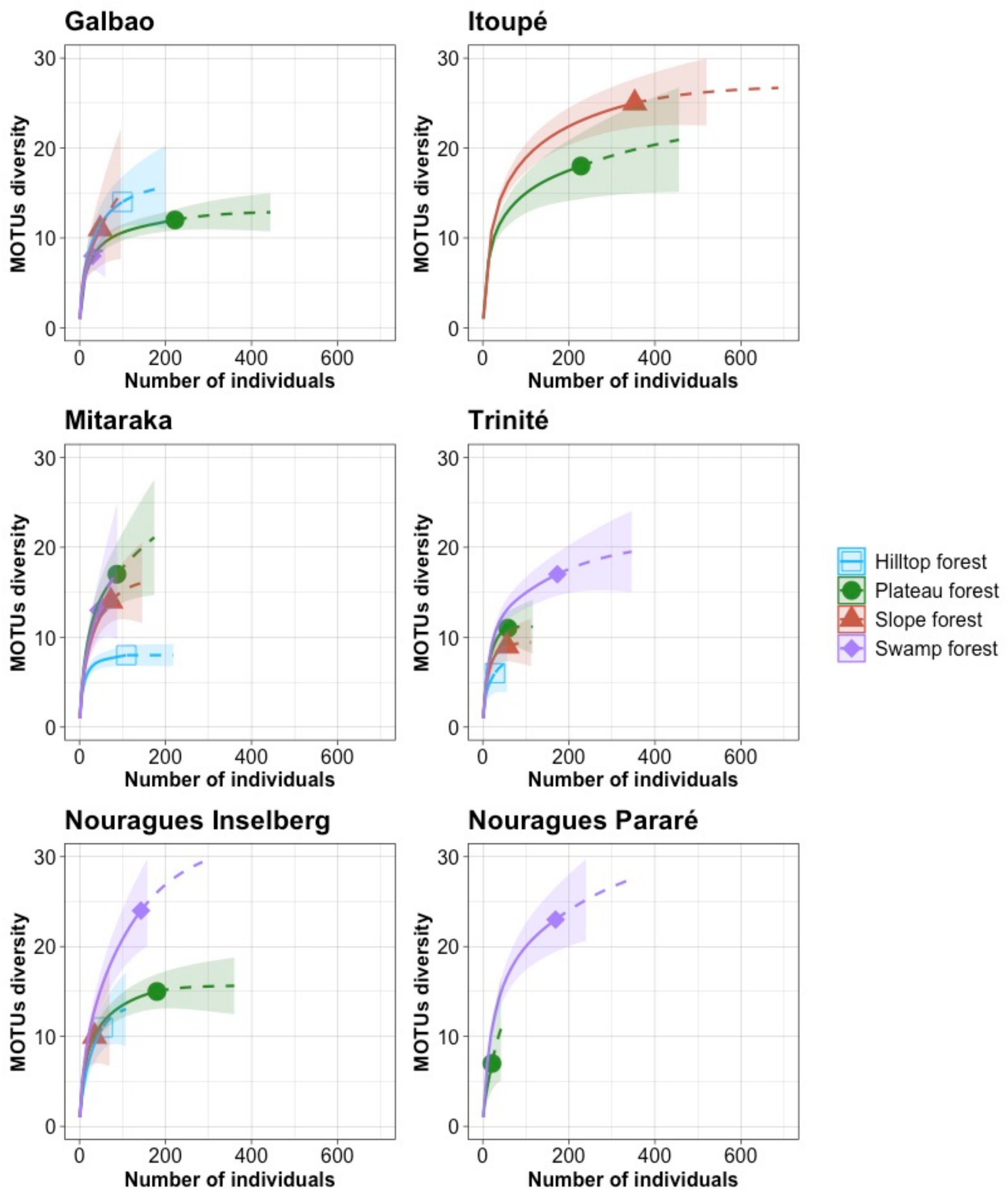


**Supplementary Figure 4:** Rarefaction and extrapolation curves showing the MOTU accumulation according to the number of sampled individuals for each main type of habitat in each of the six localities. Rarefaction curves are represented in solid lines, extrapolation curves in dashed lines; shaded areas represent a 95 % confidence intervals based on a bootstrap method with 200 replications.

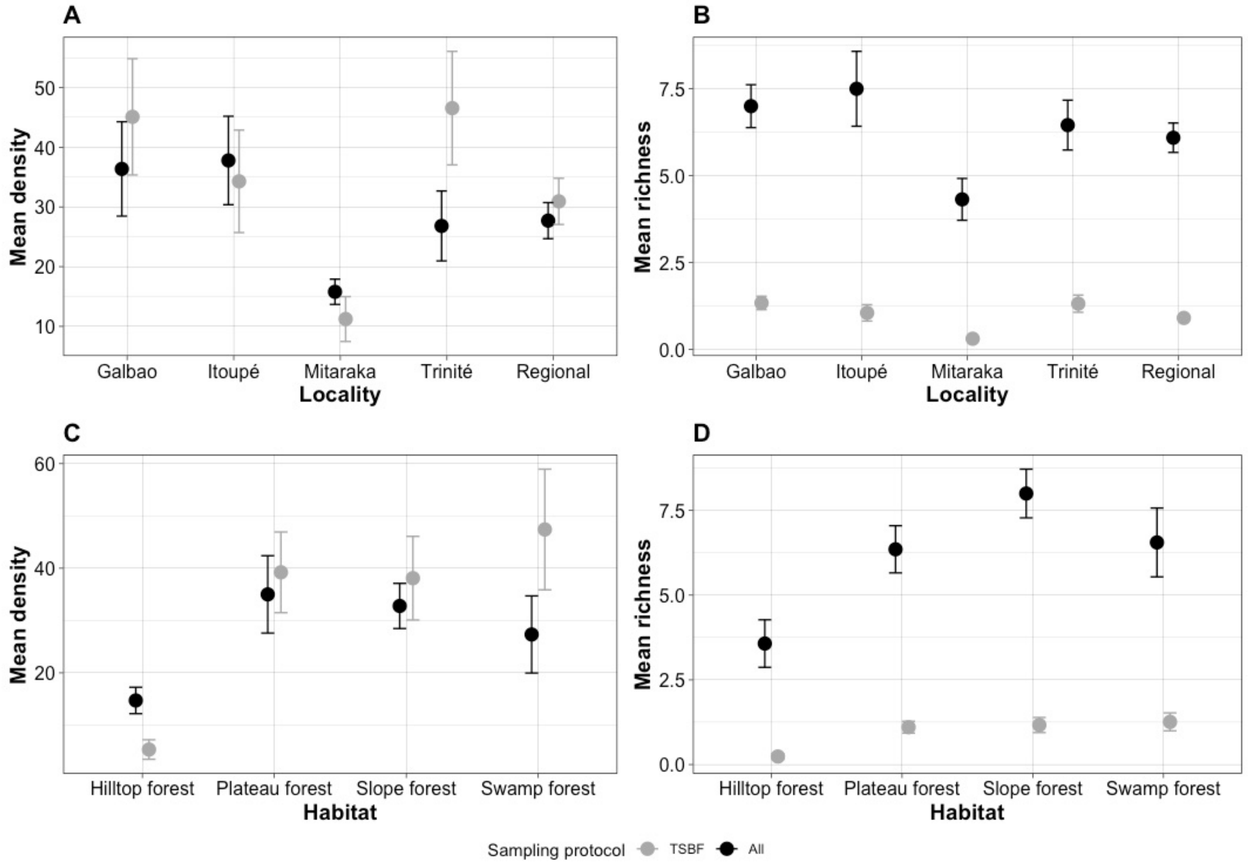


**Supplementary Figure 5:** Mean density of earthworms (A & C) expressed in earthworms per meter square for the TSBF sampling method (grey) and in total number of earthworms sampled for all sampling methods combined (black) found in a sampling plot ; and mean MOTUs richness (B & D) the number of MOTUs sampled per sampling plot. Measures are shown with the quantitative sampling (TSBF, grey) only and with the three sampling methods (all, black) ; between localities (A & B) and habitats (C & D). Points represent the mean and bars the standard error.

**Supplementary Table 1:** Sampling point with main characteristics. Sites with soil variables were used in the RDA.

| Location | Sampling point | Collecting date | Habitat | Longitude | Latitude | Elevation (m) | Clay | Fine Silt | Coarse Silt | Fine Sand | Coarse Sand | pH | Total Nitrogen (mg) | Organic Carbon (mg) | C/N | Phosphorus (mg.kg) |
|---|---|---|---|---|---|---|---|---|---|---|---|---|---|---|---|---|
| | GAL-500-1 | January 2019 | Plateau | -53.2680 | 3.6040 | 500 | 17.51 | 48.48 | 9.55 | 13.09 | 11.37 | 4.38 | 0.20 | 2.34 | 11.71 | 0.38 |
| | GAL-500-2 | January 2019 | Plateau | -53.2704 | 3.6004 | 500 | 14.23 | 58.30 | 11.13 | 13.46 | 2.88 | 4.40 | 0.15 | 1.78 | 11.95 | 0.29 |
| | GAL-500-3 | January 2019 | Plateau | -53.2662 | 3.6009 | 500 | 9.89 | 44.04 | 10.12 | 17.93 | 18.03 | 4.48 | 0.24 | 2.33 | 9.77 | 0.29 |
| | GAL-580-1 | January 2019 | Swamp | -53.2756 | 3.6016 | 600 | 5.86 | 49.22 | 14.72 | 22.17 | 8.04 | 4.09 | 0.30 | 3.57 | 11.75 | 1.07 |
| | GAL-600-2 | January 2019 | Swamp | -53.2875 | 3.6073 | 550 | 1.56 | 18.89 | 11.12 | 34.80 | 33.63 | 4.26 | 0.85 | 10.61 | 12.69 | 2.41 |
| GALBAO | GAL-700-1 | January 2019 | Hilltop | -53.2775 | 3.5964 | 700 | 9.71 | 43.66 | 13.02 | 21.33 | 12.29 | 4.24 | 0.31 | 4.48 | 14.41 | 0.78 |
| | GAL-700-2 | January 2019 | Hilltop | -53.2772 | 3.5963 | 700 | 8.60 | 41.02 | 13.69 | 21.47 | 15.24 | 3.98 | 0.36 | 4.72 | 13.10 | 0.35 |
| | GAL-700-3 | January 2019 | Hilltop | -53.2830 | 3.5886 | 700 | 11.42 | 43.25 | 10.77 | 19.64 | 14.93 | 3.88 | 0.29 | 3.96 | 13.52 | 0.89 |
| | GAL-PA1 | January 2019 | Plateau | -53.2701 | 3.6028 | 580 | | | | | | | | | | |
| | GAL-PA2 | January 2019 | Slope | -53.2756 | 3.6018 | 611 | | | | | | | | | | |
| | GAL-PA3 | January 2019 | Slope | -53.2758 | 3.5991 | 640 | | | | | | | | | | |
| | ITOU-A | January 2016 | Plateau | -53.1020 | 3.0380 | 500 | | | | | | | | | | |
| | ITOU-B | January 2016 | Plateau | -53.1080 | 3.0419 | 507 | | | | | | | | | | |
| | ITOU-C | January 2016 | Plateau | -53.1048 | 3.0393 | 440 | | | | | | | | | | |
| | ITOU-D | January 2016 | Plateau | -53.0790 | 3.0269 | 820 | | | | | | | | | | |
| | ITOU-E | January 2016 | Slope | -53.0974 | 3.0176 | 635 | | | | | | | | | | |
| | ITOU1 | January 2016 | Plateau | -53.0769 | 3.0267 | 809 | 44.76 | 25.30 | 7.47 | 6.24 | 16.23 | 4.54 | 3.66 | 4.74 | 12.96 | 2.27 |
| ITOUPE | ITOU2 | January 2016 | Plateau | -53.0834 | 3.0222 | 817 | 40.81 | 22.44 | 7.33 | 9.72 | 19.70 | 4.60 | 4.15 | 5.42 | 13.07 | 2.53 |
| | ITOU3 | January 2016 | Plateau | -53.0866 | 3.0168 | 810 | 62.44 | 8.43 | 0.57 | 3.13 | 25.44 | 4.74 | 2.77 | 3.55 | 12.87 | 1.73 |
| | ITOU4 | January 2016 | Slope | -53.0961 | 3.0333 | 578 | 78.25 | 6.92 | 1.44 | 6.74 | 6.65 | 4.54 | 2.66 | 3.24 | 12.24 | 1.27 |
| | ITOU5 | January 2016 | Slope | -53.0975 | 3.0224 | 584 | 74.91 | 8.00 | 1.31 | 6.05 | 9.73 | 4.73 | 2.72 | 3.29 | 12.13 | 1.20 |
| | ITOU6 | January 2016 | Slope | -53.0992 | 3.0115 | 585 | 79.20 | 9.85 | 1.98 | 3.01 | 5.97 | 4.61 | 2.69 | 3.37 | 12.61 | 1.53 |
| | ITOU7 | January 2016 | Slope | -53.1056 | 3.0327 | 477 | 86.85 | 3.95 | 2.49 | 3.87 | 2.83 | 4.47 | 2.24 | 2.71 | 12.11 | 1.80 |
| | ITOU8 | January 2016 | Slope | -53.1066 | 3.0221 | 439 | 14.96 | 29.22 | 19.22 | 26.34 | 10.27 | 6.22 | 2.43 | 2.22 | 9.18 | 2.00 |
| | ITOU9 | January 2016 | Slope | -53.1079 | 3.0166 | 503 | 83.21 | 7.12 | 1.52 | 3.50 | 4.66 | 5.66 | 3.06 | 2.98 | 9.77 | 1.80 |
| | MI15-A-BF | March 2015 | Swamp | -54.4650 | 2.2430 | 327 | 59.66 | 21.40 | 4.07 | 7.24 | 7.63 | 4.84 | 12.22 | 16.39 | 13.49 | 17.73 |
| | MI15-A-PEN | March 2015 | Slope | -54.4520 | 2.2380 | 344 | 55.54 | 8.00 | 0.89 | 9.87 | 25.70 | 4.52 | 4.04 | 5.32 | 13.13 | 4.60 |
| | MI15-A-PLA | March 2015 | Plateau | -54.4590 | 2.2440 | 371 | 42.24 | 8.94 | 1.02 | 8.95 | 38.84 | 4.55 | 1.92 | 2.59 | 13.41 | 2.27 |
| | MI15-BAL1 | March 2015 | Heliconia | -54.4494 | 2.2307 | 351 | | | | | | | | | | |
| | MI15-C-BF | March 2015 | Swamp | -54.4490 | 2.2380 | 310 | 34.71 | 13.34 | 5.20 | 23.08 | 23.67 | 5.12 | 3.70 | 4.27 | 11.48 | 14.07 |
| | MI15-C-PEN | March 2015 | Slope | -54.4450 | 2.2350 | 377 | 41.94 | 7.49 | 1.52 | 9.62 | 39.44 | 4.49 | 2.34 | 3.07 | 13.13 | 4.07 |
| | MI15-C-PLA | March 2015 | Plateau | -54.4440 | 2.2330 | 449 | 31.32 | 14.10 | 5.08 | 16.54 | 32.95 | 4.62 | 3.04 | 4.29 | 14.06 | 3.87 |
| | MI15-D-BF | March 2015 | Swamp | -54.4520 | 2.2330 | 318 | 34.92 | 16.07 | 7.63 | 36.44 | 4.94 | 5.02 | 3.42 | 4.05 | 11.98 | 18.53 |
| | MI15-D-PEN | March 2015 | Slope | -54.4540 | 2.2280 | 339 | 38.79 | 6.47 | 0.75 | 10.26 | 43.72 | 4.40 | 2.19 | 2.66 | 12.13 | 2.67 |
| MITARAKA | MI15-D-PLA | March 2015 | Plateau | -54.4570 | 2.2160 | 381 | 41.84 | 10.75 | 1.21 | 7.36 | 38.84 | 4.55 | 2.48 | 3.08 | 12.41 | 2.33 |
| | MI15-FS1 | March 2015 | Hilltop | -54.4673 | 2.2286 | 638 | | | | | | | | | | |
| | MI15-FS2 | March 2015 | Hilltop | -54.4369 | 2.2098 | 600 | | | | | | | | | | |
| | MI15-FTR1 | March 2015 | Transition | -54.4609 | 2.2318 | 401 | | | | | | | | | | |
| | MI15-FTR2 | March 2015 | Transition | -54.4352 | 2.2387 | 389 | | | | | | | | | | |
| | MI15-FTR3 | March 2015 | Transition | -54.4596 | 2.2331 | 395 | | | | | | | | | | |
| | MI15-SR1 | March 2015 | Rocky savannah | -54.4672 | 2.2279 | 623 | | | | | | | | | | |
| | MI15-SR2 | March 2015 | Rocky savannah | -54.4609 | 2.2329 | 427 | | | | | | | | | | |

| | | | | | | | | | | | | | | | | |
|---|---|---|---|---|---|---|---|---|---|---|---|---|---|---|---|---|
| | MI15-SR3 | March 2015 | Rocky savannah | -54.4348 | 2.2386 | 401 | | | | | | | | | | |
| | MI15-SR4 | March 2015 | Rocky savannah | -54.4389 | 2.2082 | 585 | | | | | | | | | | |
| | TRI-1 | April 2016 | Plateau | -53.4102 | 4.5982 | 144 | 39.91 | 6.43 | 3.98 | 18.29 | 31.39 | 4.26 | 1.93 | 2.38 | 12.22 | 2.00 |
| | TRI-2 | April 2016 | Plateau | -53.4146 | 4.6095 | 141 | 20.35 | 2.61 | 2.41 | 14.63 | 60.01 | 4.40 | 1.21 | 1.55 | 12.81 | 2.20 |
| | TRI-3 | April 2016 | Plateau | -53.4016 | 4.6068 | 144 | 27.19 | 3.44 | 1.25 | 12.30 | 55.82 | 4.32 | 1.41 | 1.74 | 12.28 | 2.00 |
| | TRI-4 | April 2016 | Swamp | -53.4145 | 4.5952 | 127 | 10.94 | 4.14 | 2.37 | 24.05 | 58.50 | 4.59 | 1.51 | 2.12 | 14.01 | 4.90 |
| | TRI-5 | April 2016 | Swamp | -53.4161 | 4.6030 | 121 | 21.66 | 5.15 | 2.83 | 30.87 | 39.49 | 4.46 | 1.29 | 1.37 | 10.59 | 2.07 |
| | TRI-6 | April 2016 | Swamp | -53.4120 | 4.5900 | 121 | | | | | | | | | | |
| TRINITE | TRI-8 | April 2016 | Hilltop | -53.4094 | 4.6204 | 408 | 51.66 | 15.62 | 5.79 | 22.77 | 4.16 | 4.95 | 2.85 | 2.62 | 8.99 | 3.47 |
| | TRI-CLUSIA | April 2016 | Rocky savannah | -53.4081 | 4.6187 | 390 | | | | | | | | | | |
| | TRI-CR1 | April 2016 | Stream banks | -53.4146 | 4.6029 | 124 | | | | | | | | | | |
| | TRI-PEN1 | April 2016 | Slope | -53.4116 | 4.6023 | 150 | | | | | | | | | | |
| | TRI-PEN2 | April 2016 | Slope | -53.4141 | 4.6019 | 133 | | | | | | | | | | |

**Supplementary Table 2:** Information on each of the 22 variables.

| Nature | Variable | Source | Kept for RDA analyze | Selected by *ordiR2step* |
|---|---|---|---|---|
| Spatial | Topography | This study | x | x |
| | Longitude | This study | x | x |
| | Latitude | This study | x | x |
| | Elevation | This study | x | x |
| Soil texture | Clay | This study | x | |
| | Fine silt (F.Silt) | This study | x | x |
| | Coarse silt (C.Silt) | This study | | |
| | Fine sand (F.Sand) | This study | | |
| | Coarse sand (C.Sand) | This study | x | |
| Soil chemistry | pH | This study | x | x |
| | Total nitrogen (Tot.N) | This study | x | x |
| | Organic carbon (Org.C) | This study | x | x |
| | Carbon / nitrogen ratio (C/N) | This study | x | |
| | Phosphorus | This study | x | x |
| | Bulk density | SoilGrids (Hengl et al., 2017) | | |
| | Cation exchange capacity | SoilGrids (Hengl et al., 2017) | | |
| Climatic | Annual mean Temperature | CHELSA (Karger et al., 2017) | | |
| | Temperature seasonality | CHELSA (Karger et al., 2017) | | |
| | Temperature annual range | CHELSA (Karger et al., 2017) | | |
| | Annual precipitation | CHELSA (Karger et al., 2017) | | |
| | Precipitation seasonality | CHELSA (Karger et al., 2017) | | |
